# Structure of a hibernating 100S ribosome reveals an inactive conformation of the ribosomal protein S1

**DOI:** 10.1101/382572

**Authors:** ertrand Beckert, Martin Turk, Andreas Czech, Otto Berninghausen, Roland Beckmann, Zoya Ignatova, Jürgen M. Plitzko, Daniel N. Wilson

## Abstract

To survive under conditions of stress, such as nutrient deprivation, bacterial 70S ribosomes dimerize to form hibernating 100S particles^1^. In γ-proteobacteria, such as *Escherichia coli,* 100S formation requires the ribosome modulation factor (RMF) and the hibernation promoting factor (HPF)^2-4^. Although structures of *E. coli* 100S particles have been reported^5,6^, the low resolution (18-38 Å) prevented the mechanism of ribosome inactivation and dimerization to be fully elucidated. Here we present single particle cryo-electron microscopy structures of hibernating 70S and 100S particles isolated from stationary phase *E. coli* cells at 3.0-7.9 Å resolution, respectively. Preferred orientation bias for the complete 100S particle was overcome using tilting during data collection. The structures reveal the binding sites for HPF and RMF as well as the unexpected presence of deacylated E-site tRNA and ribosomal protein S1 in the 100S particle. HPF interacts with the anticodon-stem-loop of the E-tRNA and occludes the binding site for the mRNA as well as A- and P-site tRNAs. RMF stabilizes a compact conformation of S1, which together sequester the anti-Shine-Dalgarno (SD) sequence of the 16S ribosomal RNA (rRNA), thereby inhibiting translation initiation. At the dimerization interface, S1 and S2 form intersubunit bridges with S3 and S4 and the C-terminus of S2 probes the mRNA entrance channel of the symmetry related particle, thus suggesting that only translationally inactive ribosomes are prone to dimerization. The back-to-back 100S dimerization mediated by HPF and RMF is distinct from that observed previously in Gram-positive bacteria^7-10^ and reveals a unique function for S1 in ribosome dimerization and inactivation, rather than its canonical role in facilitating translation initiation.

---

The hibernation state mediated by RMF and HPF is not only important for bacterial survival during stationary phase, but also under stress conditions, such as osmotic, heat and acid stress^1^. Since RMF and HPF are present in many clinically important bacterial pathogens, such *E. coli, Salmonella typhimurium, Yersinia pestis* and *Pseudomonas aeruginosa*^11^, the hibernation pathway represents a attractive target for development of novel antimicrobial agents. While structures of hibernating 100S particles exist, the low resolution (18-38 Å) precluded identification of the RMF and HPF binding sites^5,6^. Structures of *E. coli* HPF in complex with 70S ribosomes revealed a binding site overlapping with the A- and P-sites^12,13^, whereas *E. coli* RMF has only been visualized on a 70S ribosome from *T. thermophilus*^1^, a bacterial species that does not even contain the gene encoding RMF. Therefore, we set out to determine the structure of a physiologically relevant 100S particle isolated from nutrient deprived bacteria.

For structural analysis, *E. coli* 100S (Ec100S) particles were isolated by sucrose density gradient centrifugation of cellular lysates prepared from stationary phase cells of the *E. coli* BW25112Δ*yfi*A strain, which lacks the YfiA protein. YfiA is antagonistic to HPF, and therefore its absence enhances Ec100S formation (Extended Data Fig. 1), as reported previously^3,4^. The Ec100S sample was applied to cryo-EM grids and data was collected on a Titan Krios transmission electron microscope (see Methods). Micrograph inspection and *in silico* sorting revealed that a large proportion of the Ec100S particles dissociated into 70S ribosomes during centrifugation and/or application to the cryo-grid (Extended Data Fig. 2). Multiple attempts to reconstruct a 3D structure of the Ec100S were unsuccessful due to severe preferred orientation bias of the particles on the cryo-grid (Extended Data Fig. 2). Therefore, we focused our initial efforts on obtaining a structure of an Ec100S-derived 70S ribosome, which we termed a hibernating 70S. Subsequent 3D classification and refinement yielded a major subpopulation of hibernating 70S ribosomes that contained stoichiometric occupancy of HPF, RMF, E-site tRNA and ribosomal protein S1 (Fig. 1a-e and Extended Data Fig. 2). The final reconstruction of the hibernating 70S (Fig. 1a-e) had an average resolution of 3.05 Å (Extended Data Fig. 3a and Extended Data Table 1), with local resolution extending to 2.5 Å within the core of the ribosomal subunits (Extended Data Fig. 3b,c). Importantly, the electron densities for HPF and RMF were well-resolved (2.5-3.5 Å), enabling complete molecular models of both ligands to be built (Fig. 1f,g and Extended Data Fig. 3d-g). The E-site tRNA was less well-resolved (3.5-5.0 Å, Extended Data Fig. 3h-i), consistent with our microarray analysis revealing that multiple different tRNA species are present in the 100S particle (Extended Data Fig. 3j,k). For illustration purposes, we nevertheless fitted and refined an arbitrary deacylated tRNA into the E-site tRNA density (Fig. 1f and Extended Fig. 3i).

**Figure 1.**
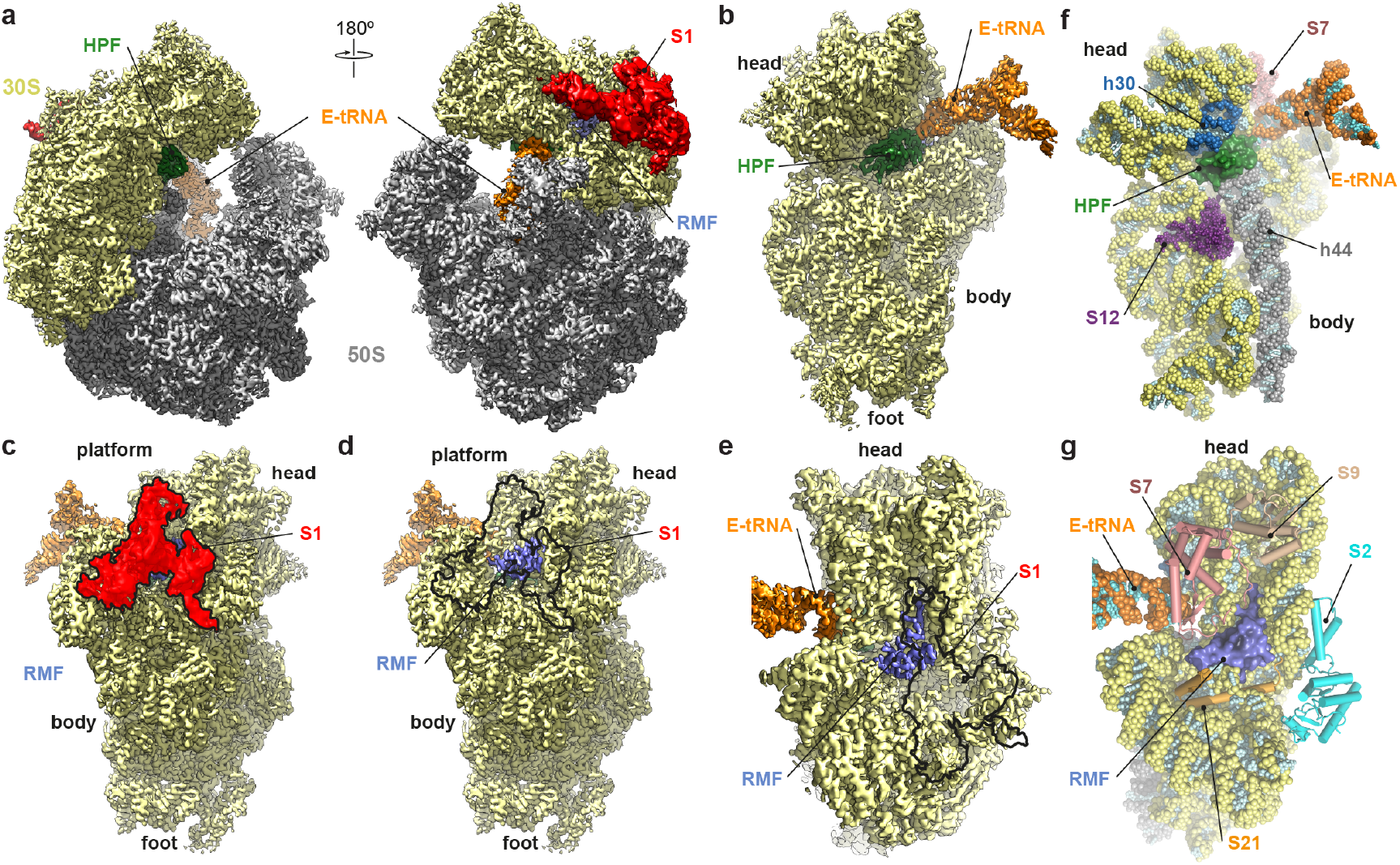
Cryo-EM structure of the hibernating 70S ribosome. **a-e**, different views of the cryo-EM map of the (**a**) hibernating 70S ribosome and (b-e) 30S subunit, highlighting the 30S (yellow) and 50S (grey) subunits and binding sites for HPF (green), RMF (blue), E-tRNA (orange) and ribosomal protein S1 (red). The black outline in (**d-e**) represents the binding position of S1 that was computationally removed to reveal the RMF binding site. f-g, Model for the 30S subunit (yellow) of the hibernating 70S ribosome focused on the binding site of (**f**) HPF (green) in the vicinity of 16S rRNA helices h30 (blue) and h44 (grey), E-tRNA (orange) and ribosomal proteins S7 (salmon) and S12 (purple), and (**g**) RMF (slate) encircled by ribosomal proteins S2 (cyan), S7 (salmon), S9 (tan) and S21 (orange).

HPF is bound within the A- and P-sites of the 30S subunit, adjacent to the E-site tRNA, where it is sandwiched between helices 30 (h30) and h44 of the 16S rRNA (Fig.1f and Fig. 2a). The location of HPF is generally similar to that observed previously^12,13^, however, the interaction with h30 differs due to head swiveling observed in the heterologous HPF-70S structure^13^ that is not observed in the hibernating 70S (Extended Data Fig. 4a-c). Moreover, HPF directly interacts with E-site tRNA (Fig. 2a,b), which was not present in the previous *in vitro* reconstituted HPF-70S structures^12,13^. Arg33 and Lys93 of HPF both come within hydrogen bonding distance of nucleotides within the anticodon-stem-loop (ASL) of the E-site tRNA (Fig. 2b), however, the nature of these interactions will depend on the exact species of the E-site tRNA. The C-terminal His95 of HPF stacks upon the nucleobase G693 within h23 of the 16S rRNA (Fig. 2b), thereby mimicking the stacking interaction observed between the first nucleotide (−3 position) of the E-site codon of a mRNA^14^ (Fig. 2c). The presence of a mixture of deacylated tRNAs in the E-site of the 100S particle (Extended Data Fig. 3j,k) is consistent with the high levels of uncharged tRNA that accumulate under conditions of nutrient deprivation^15^.This suggests that deacylated tRNA binding to the E-site may act as a signal for 100S formation, in addition to its known role in binding to the A-site of the ribosome during RelA-mediated synthesis of the alarmone ppGpp^15^.

**Figure 2.**
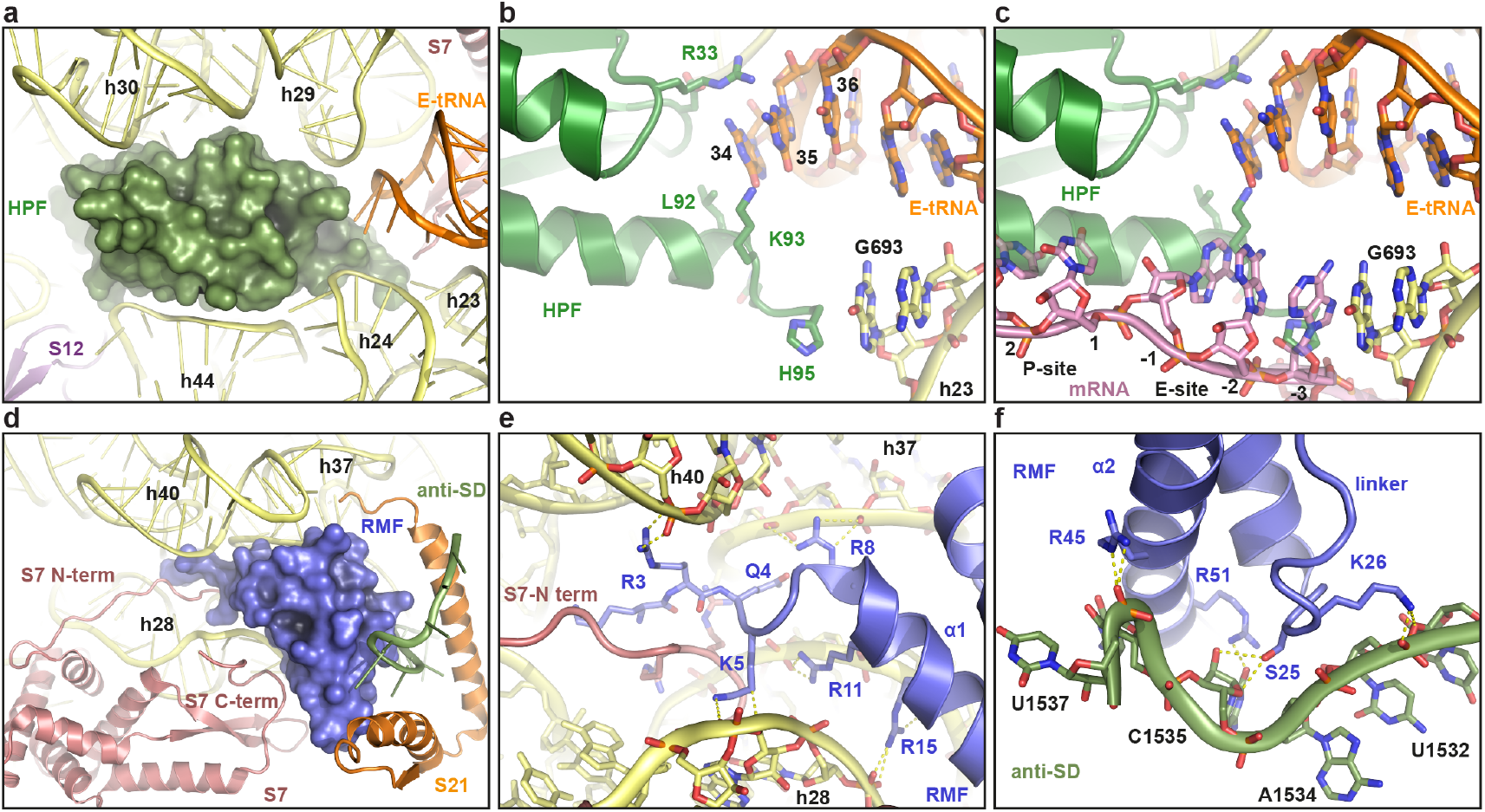
Interaction of HPF and RMF on the 30S subunit. **a**, Overview of HPF (green) binding site with interaction partners, including 16S rRNA helices (yellow), S12 (purple) and E-tRNA (orange). **b**, Contacts between HPF (green), E-tRNA (orange) and G693 of helix h23 (yellow) of the 16S rRNA. **c**, Superimposition of HPF (green) with P- and E-site mRNA (salmon, PDB ID 4V9D)^14^. **d**, Overview of RMF (blue) binding site with interaction partners, including 16S rRNA helices (yellow), S7 (salmon), S21 (orange) and anti-Shine-Dalgarno (anti-SD) of the 16S rRNA (green). e, Contacts between RMF (blue), N-terminus of S7 (salmon) and backbone of helices h28 and h40 (yellow) of the 16S rRNA. **f**, Contacts between helix a1 and a2 of RMF (blue) with nucleotides within the 3’ end of the 16S rRNA (green), including 1535-1537 of the anti-SD.

The binding site of RMF is located on the back of the 30S subunit and is better observed by computationally removing ribosomal protein S1 (Fig. 1c-e). RMF sits within a cavity created by a ring of ribosomal proteins S2, S7, S9 and S21 (Fig. 1g), with helix a1 of RMF inserting into a cleft created by 16S rRNA helices h28 on one side and h37/h40 on the other (Fig. 2d,e). RMF contains many conserved positively charged residues (Arg3, Lys5, Arg8, Arg11 and Arg15) that form a hydrogen bond network with phosphate-oxygen backbone of the 16S rRNA (Fig. 2e). Additional interactions are observed between the intertwined N-terminal extensions (NTEs) of RMF and ribosomal protein S7 (Fig. 2d,e). Helix a2 of RMF interacts with the C-terminal extension of S7 (Fig. 2d) and together with residues within the linker connecting helix a1 and a2 of RMF stabilizes a defined conformation of the 3’ end of the 16S rRNA, which encompasses part of the anti-Shine-Dalgarno (anti-SD) sequence (Fig. 2d,f and Extended Fig. 4d). The binding position of RMF and the conformation of the anti-SD observed on the hibernating 70S are incompatible with the presence of an SD-anti-SD duplex that is formed during translation initiation^16,17,14^ (Extended Data Fig. 4d-f). The position of RMF is also distinct from that observed in the heterologous RMF-70S structure^13^ (Extended Data Fig. 4g-i). In fact, the binding site for RMF observed on the *T. thermophilus* 70S does not exist on an *E. coli* 70S due to the presence of ribosomal protein S21 (Extended Data Fig. 4g-i), which is lacking in *T. thermophilus.*

The cavity in which RMF is bound is capped by a large mass of additional density that we have attributed to ribosomal protein S1 (Fig. 3a,b). Despite being the largest of the ribosomal proteins (61 kDa), S1 has not been visualized in the ribosomal X-ray structures and in some cases was even selectively removed prior to crystallization^18^. Although density for parts of S1 has been observed in cryo-EM reconstructions of various ribosomal complexes, the flexibility of S1 and/or the moderate resolution has limited the interpretation^19-22^. S1 is comprised from six structurally related oligosaccharide-oligonucleotide binding (OB)-fold domains (D1-D6), of which we observe cryo-EM density for D1 to D5 in the hibernating 70S (Fig. 3c,d and Extended data Fig. 5a,b). The N-terminal a-helix and OB-fold of D1 could be modelled based on the available crystal structure (Fig. 3e and Extended Data Fig. 5c,d) and shown to interact with ribosomal protein S2, as suggested previously^21^ (Fig. 3c). With the exception of S1-D3, which was highly flexible and poorly ordered (Extended Fig. 5b), molecular models could be unambiguously fitted for D2, D4 and D5 of S1 (Fig. 3e and Extended Fig. 5e-j). S1-D4 was particularly well-resolved (Extended Fig. 5g-h), enabling the majority of the side chains to be visualized (Extended Fig. 5k,l). The conformation of the OB-fold of S1-D4 observed here on the hibernating 70S is similar to that determined in solution by NMR^23^, whereas differences are seen for the loops connecting the β-strands (Extended Fig. 5m).

**Figure 3.**
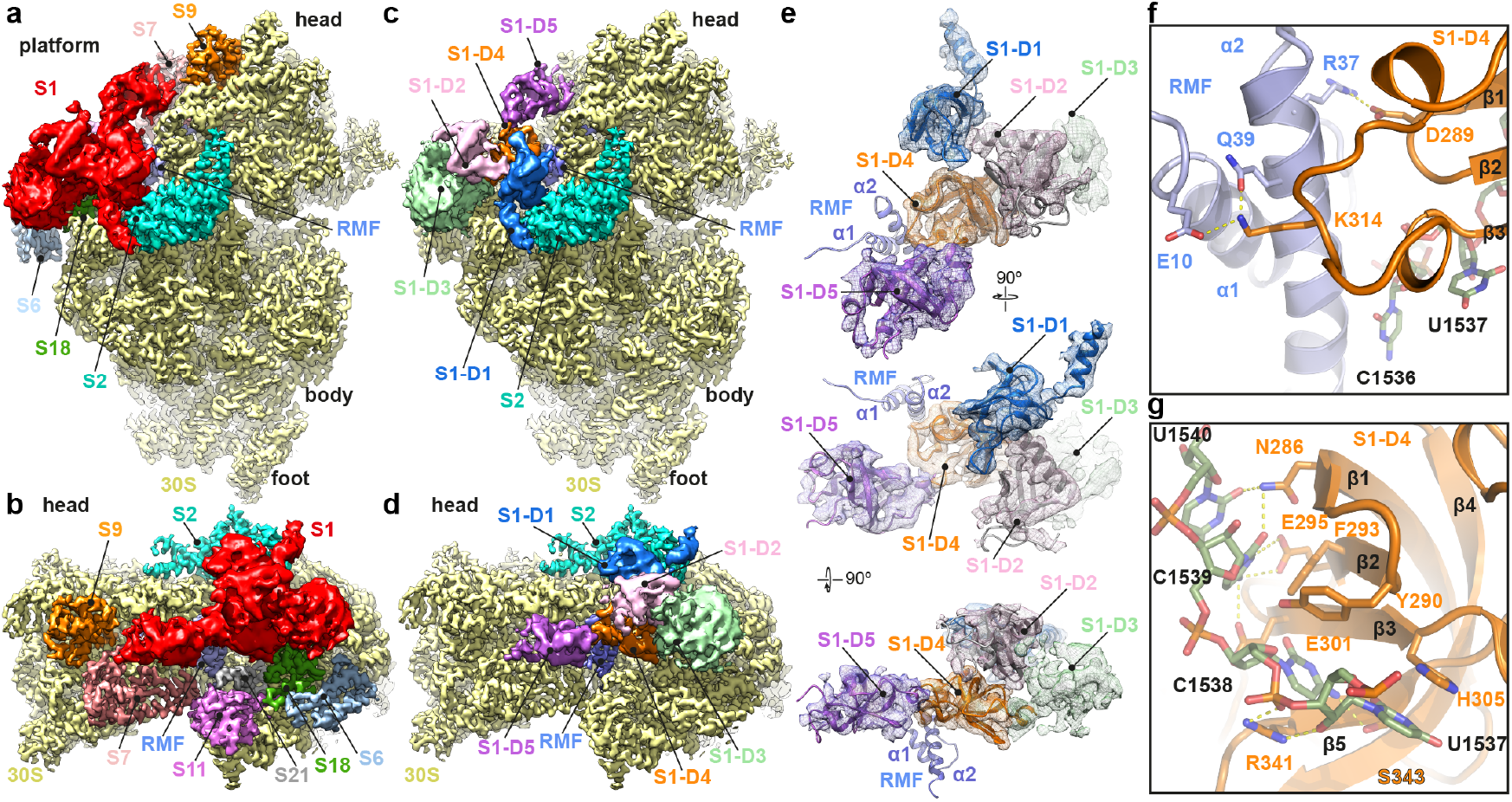
Conformation and interaction of S1 on the hibernating 70S. **a-b**, Two views of the cryo-EM map of the 30S subunit from the hibernating 70S ribosome, highlighting density for S1 (red) relative to other ribosomal proteins. **c-d**, as in (a-b), but highlighting the density for domains D1-D5 of S1. **e**, Different views of the isolated cryo-EM density (mesh) of S1 with fitted models (except D3) relative to the binding site of RMF (slate). **f**, Interaction of RMF (blue) with loops 1 and 2 of S1-D4 (orange). g, Interaction of residues within the β-sheet of S1-D4 (orange) with nucleotides of the anti-Shine-Dalgarno (anti-SD) sequence (green) at the 3’ end of the 16S rRNA.

In contrast to D1-D2 of S1 that are involved in ribosome binding^24,21^, D3-D6 have been shown to interact with mRNA^24^. S1 facilitates unwinding of mRNA secondary structures and promotes placement of the start codon of the mRNA within the ribosomal decoding site to allow translation initiation to occur^24-26^. Biophysical studies have indicated that S1 has a very elongated shape (about 230 Å long) in solution, as well as on the ribosome, consistent with the idea that while the N-terminal domains (NTDs) of S1 anchor it to the ribosome, the C-terminal domains (CTDs) extend into the cytoplasm to fish for mRNAs^24^. This would explain why the S1-CTDs are poorly resolved in the available cryo-EM structures^19-21^, except for an elongated S1 conformation observed upon interaction with the RNA polymerase^22^ (Extended Data Fig. 6a-c). By contrast, we observe a very compacted conformation (100 Å for the longest edge) for S1 on the ribosome with the S1-CTDs folding back onto the 30S subunit, rather than extending into the solvent (Fig. 3c-e and Extended Data Fig. 6d). Specifically, S1-D4 folds back under S1-D2 into the ribosomal cavity where RMF is bound, thereby directing S1-D5 towards the head of the 30S subunit (Fig. 3c,d). The compacted conformation of S1 is stabilized by the direct interactions observed between S1-D4 and RMF (Fig. 3f) as well as with the anti-SD sequence of the 16S rRNA (Fig. 3g). Two flexible loops, located between strands β1-β2 and β3-β4 of S1-D4 contact RMF. Potential hydrogen bonds are observed between Asp289 in the β1-β2 loop of S1-D4 and Arg37 of RMF, as well as between Lys314 in β3-β4 of S1-D4 and Glu10 and Gln39 in helix a1 and a2, respectively, of RMF (Fig 3f). Multiple contacts are established between residues located in the β-sheet surface formed by strands β1-β3 of S1-D4 and nucleotides C1537-U1540 of the anti-SD sequence (Fig. 3g and Extended Data Fig. 5l). These include stacking interactions between His305 and Phe293 of S1-D4 with the bases of nucleotides U1537 and C1539, respectively, as well as potential hydrogen bond interactions between Asn286, Glu295 and Glu301 of S1-D4 with 16S rRNA nucleotides C1539 and U1540 (Fig. 3g). In addition, Arg342 and Ser343 within the β5 strand of S1-D4 are within hydrogen bonding distance of nucleotides U1537 and C1538 (Fig. 3g). The surface of S1-D4 seen here to interact with the anti-SD of the 16S rRNA is the same as that shown by NMR to interact with mRNA in solution^27^, suggesting that the compact conformation adopted by S1 is an inactive form that is incompatible with its canonical role during translation initiation.

To assess whether the results revealed by the structure of the hibernating 70S reflect the physiological state of the hibernating 100S, we renewed our attempts to determine a structure of the complete hibernating 100S particle. To overcome the preferred specimen orientation bias of the 100S particles on the cryo-grids, we employed single particle cryo-EM using tilting (40°) during data collection^28^. In contrast to the severe anisotropy displayed by the 0° untilted data, the 40° tilt data showed a significant improvement in the distribution of Euler angles (Extended Data Fig. 7a-c). Next, the relative orientation and distance between 70S particles on each image was determined and only 100S particles were selected that contained a C2 symmetry related 30S-30S back-to-back arrangement (Expanded Data Fig. 7d-f). 3D classification was performed (Extended Data Fig. 8), producing multiple classes that exhibited a high degree of movement (up to 8° tilting) between the 70S ribosomes of the 100S particle (Extended Data Fig. 9a). Therefore, the most stable classes were combined and refined, yielding a structure of the hibernating 100S (Fig. 4a) with an average resolution of 7.9 Å (Extended Figure 9b). Remarkably, a complete molecular model could be generated for the hibernating 100S including HPF, RMF, E-tRNA and S1 (Fig. 4b) by a rigid body fit of the intact structure of the hibernating 70S into the density for each of the symmetry-related 70S ribosomes of the 100S particle (Extended Data Fig. 9c-e and Supplementary Video 1). Thus, the molecular details revealed by the high-resolution structure of the hibernating 70S ribosome are valid for the complete 100S particle, at least within the limits of the resolution. In addition, extra density is observed in the 100S particle for S1-D6, which extends from the head of one 70S ribosome towards the head of the symmetry related 70S particle where it contacts S10 (Fig. 4c and Extended Data Fig. 9f-g).

**Figure 4.**
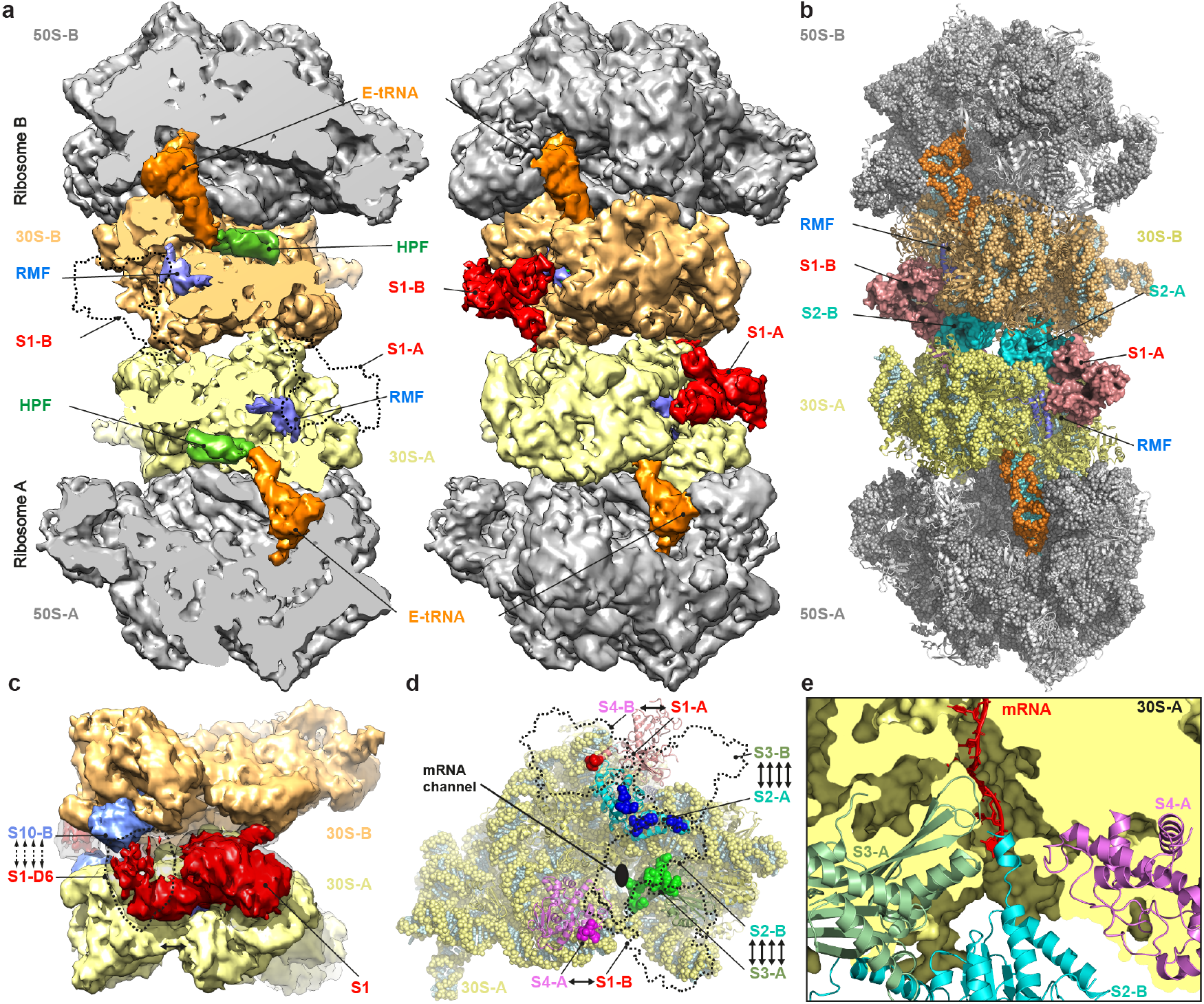
Cryo-EM structure of the hibernating 100S particle. **a**, Transverse section and overview of the cryo-EM structure of the hibernating100S particle, highlighting the binding positions of HPF (green), RMF (blue), E-tRNA (orange) and ribosomal protein S1 (red) on the 30S (30S-A, yellow and 30S-B, tan) and 50S (grey) subunits of ribosome A and B. The dashed line represents the binding position of S1 that was computationally removed to reveal the RMF binding site. **b**, Molecular model of the 100S hibernating ribosome generated from rigid body fitting of the structure of the hibernating 70S ribosome in the cryo-EM (see also Extended Data Fig. 9e). **c**, Views onto the dimerization interface of the hibernating 100S showing the respective interaction of S1-D6 (red) with ribosomal protein S10 (blue). **d**, View onto the 30S-A subunit showing the intersubunits bridges (indicated by spheres) between S1-A and S4-B (and S1-B with S4-A), as well as between S2-A and S3-B (S2-B and S3-A). The dashed lines indicate the respective S1-S4 proteins of the 30S-B subunit. The black circle indicates the position of the mRNA entrance channel on the 30S-A subunit. e, Transverse section of the mRNA entrance channel of the 30S-A subunit illustrating the interaction of the C-terminal helix of S2-B (cyan), which interacts with S3-A (green) and S4-A (magenta). The position of mRNA (red) is indicated for reference.

On the basis of the heterologous RMF-70S complex structure^13^, dimerization was proposed to be mediated by an RMF-induced reorientation of the 30S head, which increased the contact surface between the head domains of the respective subunits in the 100S particle. By contrast, we observe no head swiveling in the structures of the hibernating 70S and 100S (Extended Data Fig. 10a-d), and also no contact between the head domains of the respective subunits in the 100S particle (Fig. 4a,b). Instead, an analysis of the dimerization interface between the two 30S subunits in the 100S particles reveals the presence of two major intersubunit bridges where S1 and S2 from each 30S interact with S4 and S3, respectively, of the symmetry related 30S (Fig. 4d). While the N-terminus of S1 forms a relatively defined contact with a loop region (residues 24-26) near the N-terminus of S4, a much larger surface area is buried by the interaction between S2 and S3 (Fig. 4d). Additionally, the C-terminus of S2 from one 30S extends towards the mRNA entrance channel of the symmetry related 30S. In the hibernating 70S, density for the C-terminal a-helix of S2 terminates at residue 225, whereas interaction with the surrounding positively charged residues of S3 and S4 in the hibernating 100S stabilizes an additional eight residues (226-233) at the end of the a-helix (Fig. 4e and Extended Data Fig. 11). Without even considering the further eight residues (234-241) at the very C-terminus of S2 that are not observed in the structure, the C-terminal a-helix of S2 effectively plugs the mRNA entrance channel, suggesting that such an interaction would not be possible on a translating ribosome containing full-length mRNA (Fig. 4e). The probing of a vacant mRNA entrance channel by S2 appears analogous to the C-termini of ribosome rescues factors, such as SmpB, ArfA and ArfB^29^, suggesting that only translationally inactive ribosomes are prone to dimerization (Fig. 4e).

In conclusion, the structure of the hibernating 100S from Gram-negative bacteria reveals that RMF and HPF mediate 70S dimerization indirectly via stabilization of ribosomal proteins S1 and S2 (Extended Data Fig. 11), which is structurally and mechanistically unrelated to 100S formation observed in Gram-positive bacteria where the long form of HPF directly participates in the dimerization interface^7-10^ (Extended Data Fig. 10e-g). While ribosomal protein S1 is well-known for its role during translation initiation, trans-translation and Qβ replication^24,30,26^, here we now expand the functional role of S1 to also include ribosome inactivation and 70S dimerization.

## Acknowledgements

We thank Susanne Rieder and Charlotte Ungewickell for expert technical assistance. This research was supported by grants from the Deutsche Forschungsgemeinschaft SPP-1879 (to D.N.W), CZ234/1-1 (to A.C.), FOR1805 (to D.N.W., Z.I. and R.B.).

## Author Contributions

D.N.W. designed the study. B.B. prepared the cryo-EM sample. O.B. collected the single particle cryo-EM data, which was processed by B.B. The tilt data was collected and processed by M.T. A.C. performed the microarray analysis. B.B. built and refined the molecular models and generated the figures. B.B., M.T., R.B., Z.I., J.P. and D.N.W. interpreted the results. B.B. and D.N.W. wrote the paper.

## Author Information

Reprints and permissions information is available at xxx. The authors declare no competing financial interests. Readers are welcome to comment on the online version of the paper. Correspondence and requests for materials should be addressed to D.N.W.

## METHODS

### 100S ribosome preparation

The *E. coli* BW25112Δ*yfiA* strain was first grown for 33 h in M9 minimum salt media (33.7 mM Na2HPO4, 22 mM KH2PO4, 8.55 mM NaCl, 9.35 mM NH4Cl, 1 mM MgSO4, 0.3 mM CaCl2) complemented with 0.4 % D-glucose. After collecting cells at 5000 x g at 4°C for 15 min, the cells were re-suspended in B100S buffer (25 mM Hepes pH 7.5, 100 mM KOAc, 15 mM Mg(OAc)2, 1 mM dithiothreitol) and lysed 3 times at 15,000 psi in an ‘microfluidics model 110I lab homogenizer’. The lysate was cleared at 12,000 x g at 4°C for 5 min and ribosomes were pelleted through a 25% (w/v) sucrose cushion (in B100S buffer supplemented with 0.01% n-dodecyl D-Maltoside) for 16 h by centrifugation at 57,690 x g (using Ti70 rotor). The ribosomes were re-suspended in B100S buffer and the 100S ribosomes were purified by 10-40% linear sucrose gradient (centrifugation at 89,454 x g for 4 h in a SW-28 rotor). The 100S fractions were collected using a gradient station (Biocomp) equipped with an Econo UV Monitor (Biorad). The purified 100S ribosomes were concentrated by centrifugation at 53,782 x g for 3h (in a Ti70.1 rotor) and re-suspended in B100S buffer to a final concentration of 450 OD_260_/mL.

### tRNA identification by microarray

RNA extracted from the 100S ribosome fraction was ligated to a Cy3-labeled RNA/DNA hybrid oligonucleotide with T4 DNA Ligase (Thermo Scientific) in ligation buffer containing 15% (v/v) DMSO for 16 h at 16°C. RNA was purified by phenol:chloroform extraction and ligation efficiency verified on denaturing 10% polyacrylamide gel electrophoresis. Labeled tRNA samples were loaded on a microarray containing 24 replicates of full-length tDNA probes recognizing 40 *E. coli* tRNA isoacceptors and hybridized for 16 h at 60°C. Fluorescence signals of microarrays were recorded with a GenePix 4200A scanner (Molecular Devices) and analyzed with the GenePix Pro 7.0 software.

### Negative-stain electron microscopy

100S ribosome complexes were diluted in B100S buffer to final concentrations of 4-5 OD_260_/ml in order to determine the optimal ribosome density for grid preparation. The specimens were stained with uranyl acetate and micrographs were collected using a Morgagni transmission electron microscope (FEI, 80 kV, wide angle 1K CCD at direct magnifications of 72K.

### Single particle cryo-EM reconstruction of the hibernating 70S ribosome

A total of 4 OD_260_/ml of *E. coli* 100S sample were applied to 2 nm pre-coated Quantifoil R2/2 UltrAuFoil holey gold grids and vitrified using Vitrobot Mark IV (FEI). Data collection was performed using EM-TOOLS (TVIPS GmbH) on a Titan Krios TEM equipped with a Falcon II direct electron detector (FEI) at 300 kV at a defocus range between 0.7-3.5 μm. All micrographs were recorded under low dose conditions and fractionated into 10 frames (with a dose per frame of 2.5 e^-^/Å^2^) with a pixel size of 1.084 Å. Dose fractionated movies were aligned using Unblur^31^. Determination of the CTF, defocus values, and astigmatism were performed using the GCTF^32^. Micrographs showing Thon rings beyond 3.8 Å were manually inspected for good areas and power-xspectra quality. Automatic particle picking was then performed using Gautomatch (http://www.mrclmb.cam.ac.uk/kzhang/) and single particles were processed using Relion 2.1^33^. The initial set of picked particles was first subjected to an extensive 2D classification resulting in 625,839 particles. 3D refinement was performed using *E. coli* 70S ribosome as a reference structure (Extended Figure S2a). The 100S-derived 70S particles (Extended Figure S2b) were then further 3D classified (Extended Figure S2c), resulting in a major population of 188,304 particles, yielding after a final round of 3D refinement a reconstruction with an average resolution of 3.0 Å according to Relion gold standard and FSC0.143 criterion^34,35^ (Extended Figure S2d). The final maps were sharpened by dividing the maps by the modulation transfer function of the detector and by applying an automatically determined negative B factor to the maps using Relion 2.1^33^. All the maps were filtered according to local resolution using SPHIRE^36^. Local resolution of the final maps was calculated using ResMap^37^. Cross-validation against over-fitting was performed as described^38^ and the statistics of the refined model were obtained using MolProbity^39^ (Extended Data Table 1).

### Data collection of 40° tilted images

The *E. coli* 100S sample at a final concentration of 4.5 OD260/ml was applied to freshly glow-discharged Quantifoil R3.5/1 grids pre-coated with a 2 nm thick carbon support. Grids were vitrified on a Vitrobot mark IV (FEI) after 3.0-3.5 sec blotting time to remove excess liquid. Data collection was performed on a Titan Krios TEM (FEI) operating at 300 kV equipped with a GIF energy filter with the slit width set to 20 eV. Micrographs were recorded at a 40° stage tilt^28^ with a K2 direct electron detector at a nominal magnification of 105k (pixel size of 1.35 Å) and a defocus range from 1.2-3.5 μm. A total of 5,570 micrographs was collected in dose-fractionation mode with a cumulative dose of 51 e^-^/Å^2^ distributed over 48 frames. Frame-alignment and dose weighting were done with MotionCor2^40^. Defocus parameters were estimated using GCTF v1.06^32^. After micrograph quality inspection 5,165 were selected for further processing. Particle picking was done with Gautomatch using projections from a 70S reference (EMD 8237).

### Data-collection of untilted images

The 100S samples were diluted to a final concentration of 18 OD260/ml and applied to freshly glow-discharged holey carbon Quantifoil R2/1 grids without an additional thin layer of carbon. Vitrification and data collection were performed similar to the tilted data with the following differences: A different Titan Krios was used, therefore the calibrated pixel size on the K2 camera at 105k magnification was slightly different at 1.34 Å. During data collection the stage tilt was 0° and the cumulative dose of 38 e^-^/Å^2^ was distributed over 33 frames. A total of 1,882 micrographs were collected. Frame-alignment and dose weighting were done with MotionCor2^40^. Defocus parameters were estimated using CTFFIND3^41^. After micrograph quality inspection 1,278 were selected for further processing. Particle picking was done with Gautomatch using projections from a 70S reference (EMD 8237) and particle coordinates were imported into Relion 2.1^33^.

### *In silico* selection of 100S particles based on 70S orientation assignment

Selection of 100S particles was based on the observation from previous cryo-EM reconstructions^5,6^ that the 100S was comprised of two 70S ribosomes oriented with a 30S-30S back-to-back orientation (Extended Data Figure S7e). The 70S particles satisfying both distance and orientation constraints were selected as 100S pairs in the following way: First, relative orientations of the picked 70S particles were obtained from one round of projection matching to a 70S reference. Second, for each particle in a micrograph the orientations obtained were compared to all other particles in its vicinity (Extended Data Figure S7f). By setting an upper and lower limit to the radius from 180 Å to 280 Å for the distance constraint, the search area was constrained to approximately one ribosome around the particle of interest. For particles falling within the search range, difference angles for each pair of particles were calculated (i.e. the angle needed to rotate and superimpose particle A onto particle B). Particles oriented towards each other with their 30S subunits on the same axis would have an expected difference angle around 180°. To avoid excluding potential off-axis pairs during the selection process, the tolerance for the difference angle was loosely set to ±60°. To select for particles oriented towards each other with their 30S subunits, a vector pointing from the centre of the particle towards the 30S subunit was introduced into the reference (Extended Data Figure 7d-f). The vector was rotated together with the particle and particle pairs that were pointing with the newly introduced vector to each other were considered further. It was necessary to take the particle coordinates (x,y) as well as the orientations into account to distinguish between particles facing each other with 30S-to-30S or 50S-to-50S orientations. Finally, the centre between the two 70S particles (_rlnOffset corrected) satisfying all the above criteria was used as the centre for the newly selected 100S particles.

### Single particle reconstruction of the 100S

After the *in silico* 100S particle selection 150,000 particles were obtained. By collecting the data at 40° tilt a defocus gradient of more than 300 nm in each micrograph was introduced which required defocus estimation on a per-particle basis^28^. A refinement of the defocus values taking into account particle coordinates was done using GCTF v1.06^32^. The obtained particle coordinates were then imported into Relion 2.1^33^ and their defocus values updated according to the GCTF per-particle estimates^28^. Particles were then subjected to 50 rounds of 3D classification into 8 classes using a mask around the full 100S particle. Particles from the three classes where both 70S ribosomes were well resolved were selected yielding 58,000 particles. To further select for the most homogenous subset of 100S particles a second round of 3D classification was performed with the same mask as in the first 3D classification run and by increasing the number of classes to 12 and rounds to 100. Selecting only a subset of 100S classes which were similar in the slices through their mid-section resulted in 18,000 particles which refined to 7.9 Å (Extended Figure 8c,9b-c). Both 70S-A and 70-B particles comprising the 100S appeared very similar. Difference maps between both 70S halves revealed no additional density, allowing for applying C2 symmetry to the refinement, which resulted in a 100S map at 7.2 Å (Extended Figure 8c).

### Molecular model of the hibernating 70S and 100S particles

The molecular model for the ribosomal proteins and rRNA core was based on the molecular model from the recent cryo-EM reconstruction of the *E. coli* 70S ribosome (PDB ID 5O2R)^42^. The models were rigid body fitted into the cryo-EM density map using UCSF Chimera^43^ followed by refinement using Coot^44^. For *E. coli* HPF and RMF, the molecular models from HPF and RMF in complex with the *T. thermophilus* 70S ribosome (PDB IDs 4V8G/H)^13^ was extracted and fitted into the density followed by manual refinement in Coot. The initial template for the E-site tRNA was deacylated tRNA^Val^ taken from the ErmBL-70S (PDB ID 5JTE)^45^ and fitted to the density using Chimera and refined in Coot. For the S1-D1, the molecular model was based on the crystal structure of *E. coli* ribosomal protein S2 in complex with N-terminal domain of S1 (PDB ID 4TOI)^21^, whereas S1-D2 was built using the NMR structure of *E. coli* S1-D2 (PDB ID 2MFL)^46^, which was fit to the density as a rigid body. The initial model for S1-D4 was built using the NMR structure of S1-D4 (PDB ID 2KHI)^23^ and then manually adjusted and refined in Coot. S1-D5 was built based on S1-D4 structure and fitted into the density as a rigid body. All atomic coordinates (excepted the S1-D1, -D2 and -D5) were refined using *phenix.real_space_refine,* with restraints obtained by *phenix.secondary_structure_restraints^47^*.

## Figure preparation

Figures showing electron densities and atomic models were generated using either UCSF Chimera or PyMol Molecular Graphic Systems (Version 1.8 Schrödinger).

## Extended Data Figures

**Extended Data Figure 1.**
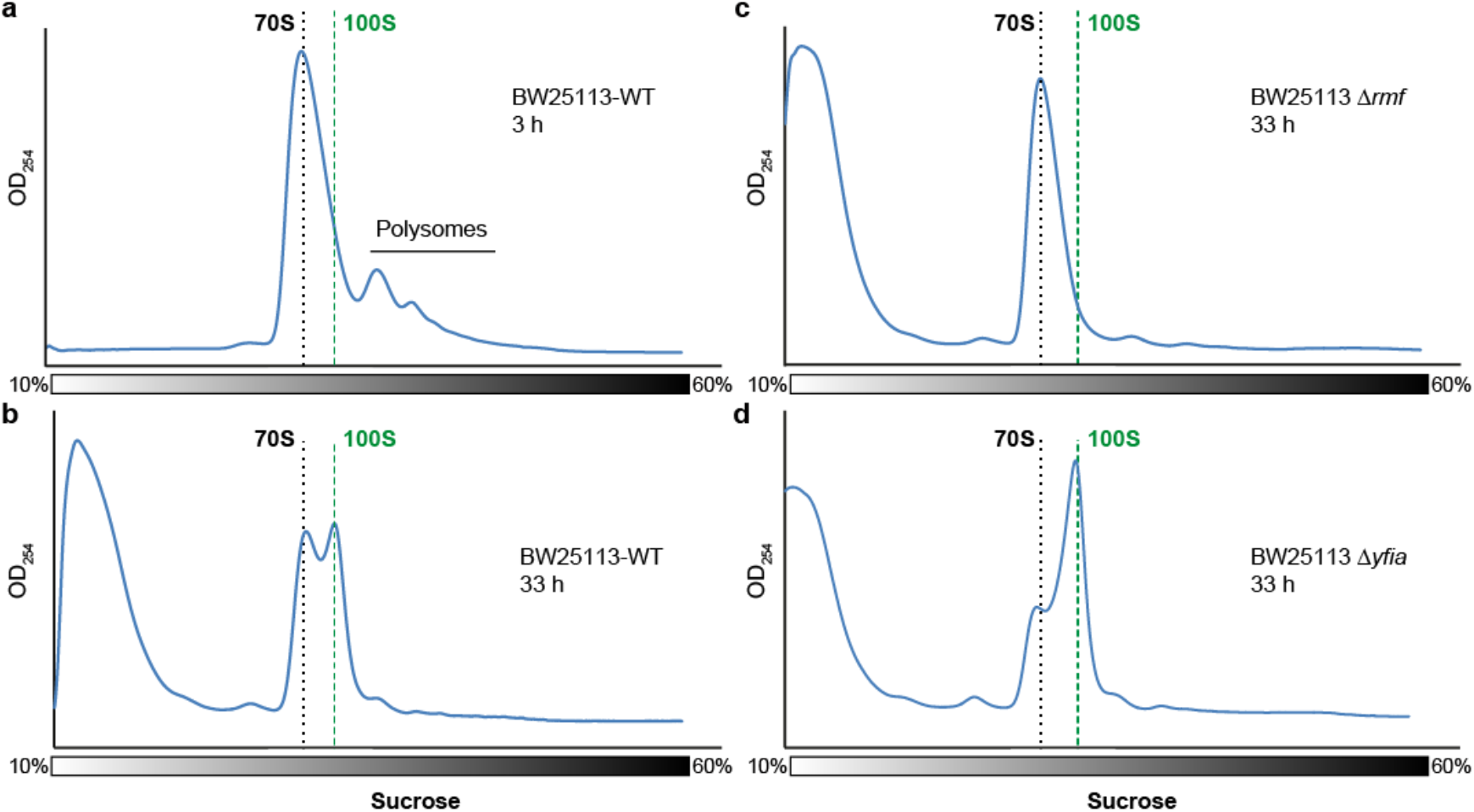
Purification of *E. coli* 100S particles. **a-d**, Sucrose density (10-60%) gradient centrifugation profiles (monitored with an optical density of 254 nm; OD254) of lysates prepared from (**a-b**) wildtype *E. coli* BW25113 strain in (**a**) exponential phase (after 3 hrs) and (**b**) stationary phase (after 33 hrs), (**c-d**) stationary phase (after 33 hrs) from (**c**) BW25113 *Drmf* strain lacking the gene for RMF, and (**d**) BW25113 *DyfiA* strain lacking the gene for YfiA. The dashed black and green lines indicate the position of 70S and 100S particles, respectively, and the position of polysomes in (a) is indicated.

**Extended Data Figure 2.**
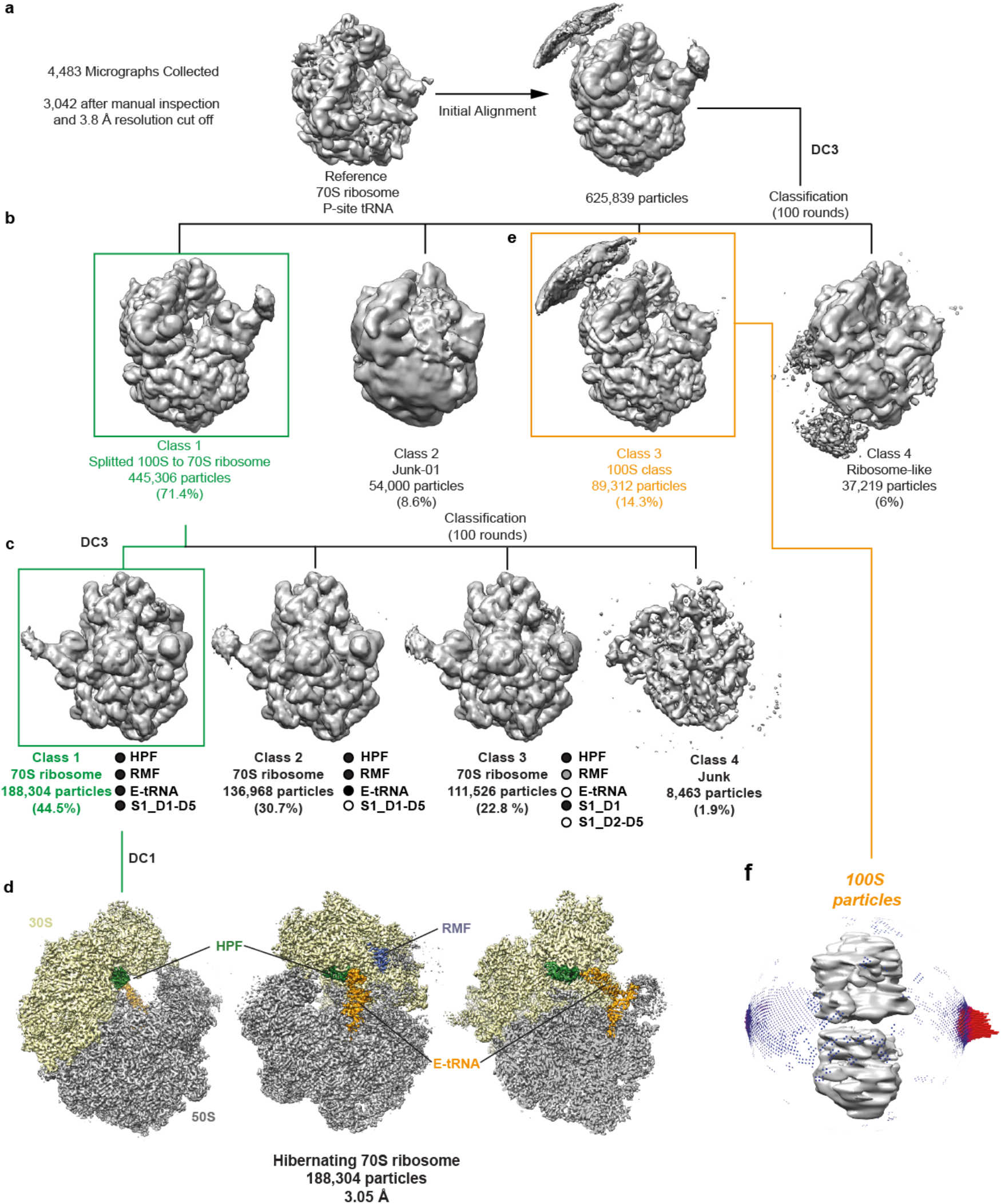
*In silico* sorting of the hibernating 70S ribosome. **a**, The complete dataset was initially aligned against an P-site tRNA containing *E. coli* 70S ribosome, yielding 625,839 particles (assigned as 100%). **b**, Following 3D classification for 100 rounds in Relion^33^ four classes were generated containing 100S-derived 70S ribosomes (class 1, 445,306 particles, 71.4%), junk particles (class 2, 54,000 particles, 8.6%), 100S particles (class 3, 89,312 particles, 14.3%) and poorly-aligning ribosomal particles (class 4, 37,219 particles, 6%). **c**, 445,263 particles from class 1 were sorted further and the resulting class 1 (188,304 particles, 44.5%) containing stoichiometric occupancies of HPF, RMF, E-tRNA and S1_D1-D5 was refined to (d) yield a final reconstruction with an average resolution of 3.05 Å (0.143 FSC) average resolution. **e**, Preferred orientation bias of the 100S particles in class 2 from (b) prohibited further refinement of this class.

**Extended Data Figure 3.**
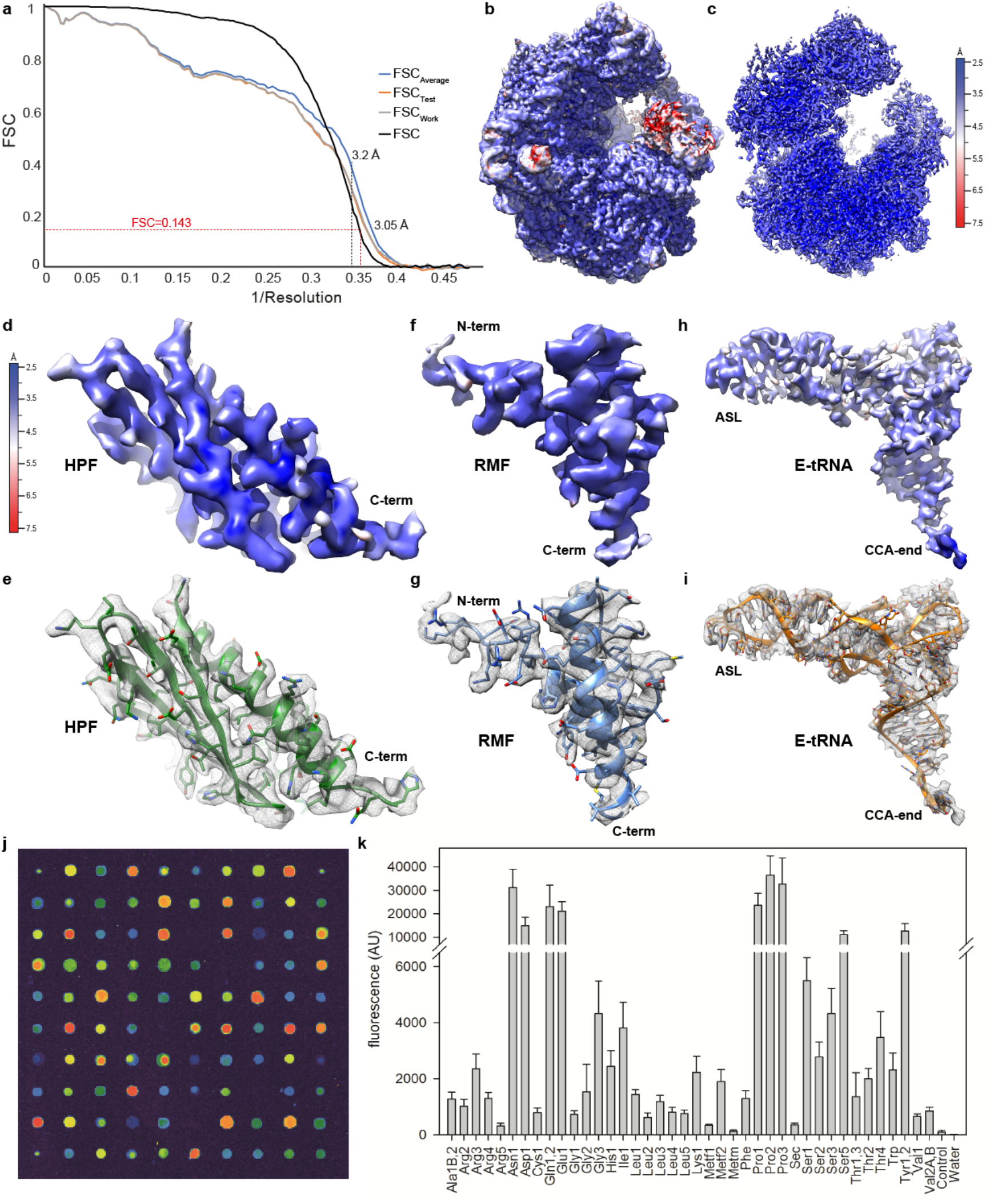
Resolution of the hibernating 70S ribosome. **a**, Fourier-shell correlation (FSC) curve (black) of the final refined map, indicating that the average resolution of the hibernating 70S ribosome is 3.05 Å (at FSC 0.143), as well as FSCaverage (blue) with self- and cross-validated correlations FSCwork (grey) and FSCtest (red), respectively. **b**, Overview and (**c**) transverse section of the final refined cryo-EM map of the hibernating 70S ribosome colored according to local resolution. **d-i**, Isolated cryo-EM density for (**d-e**) HPF, (**f-g**) RMF and (**h-i**) E-tRNA, (**d,f,h**) colored according to the local resolution or (**e,g,i**) represented as mesh (grey) with fitted molecular models. The N- and C-termini of HPF and RMF as well as the anticodon-stem-loop (ASL) and CCA-end of the tRNA are indicated. (**j**) One representative block out of 12 from the microarray analysis of tRNAs present in the *E. coli* 100S sample. (**k**) Microarray quantitation of results from all 12 microarray blocks.

**Extended Data Figure 4.**
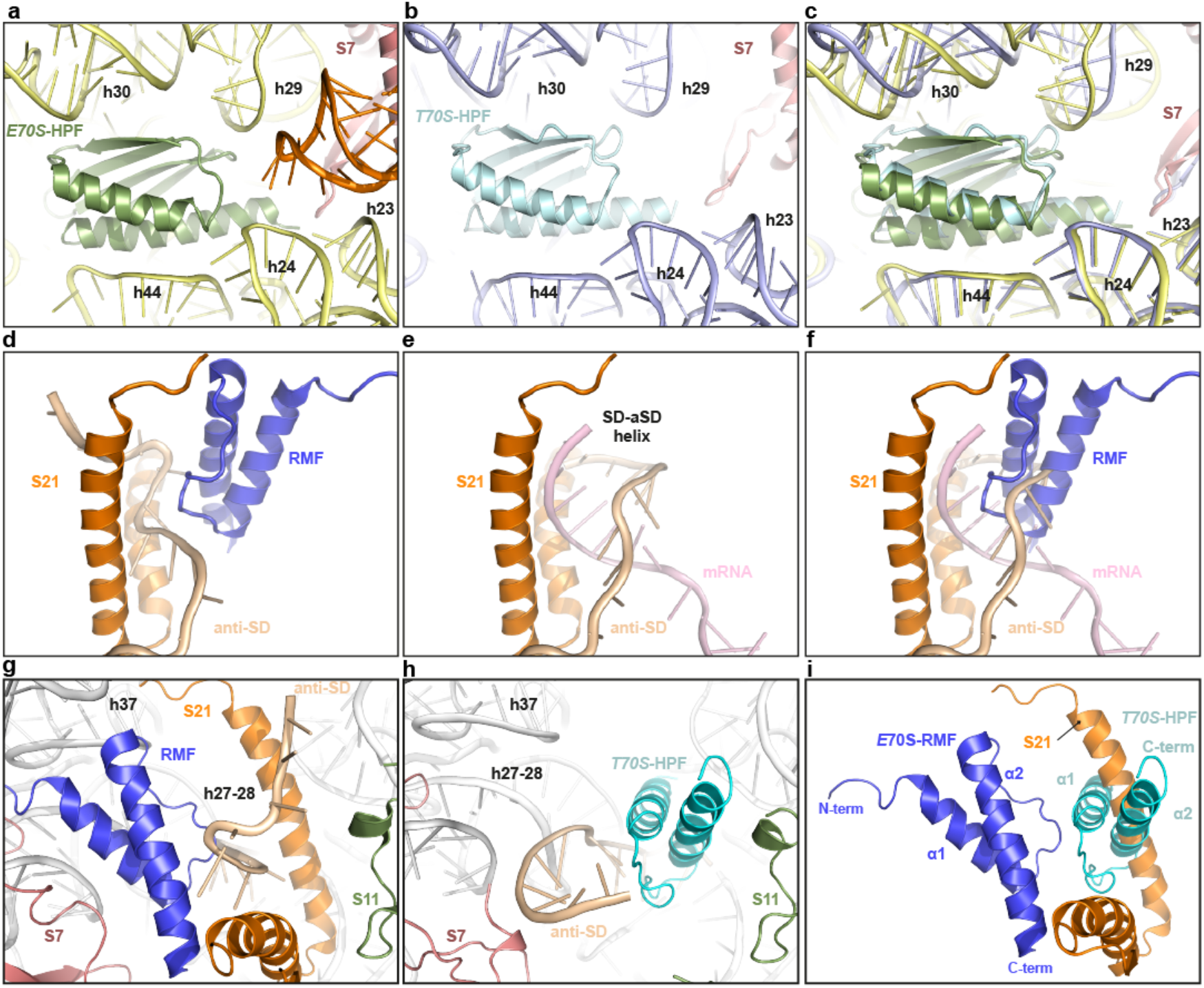
Interaction of HPF and RMF on the hibernating 70S. **a**, Binding site of *E. coli* HPF (green) on the hibernating 70S (E70S-HPF) compared with the (**b**) binding site *E. coli* HPF (cyan) on the *T. thermophilus* 70S (T70S-HPF), and (**c**) superimposition of (**a**) and (**b**). Helices of the 16S rRNA (yellow), ribosomal protein S7 (salmon) and E-tRNA (orange) are shown for reference. **d**, Binding site of *E. coli* RMF (blue) on the hibernating 70S with S21 (orange) and anti-SD (tan). e, Binding site of the helix formed by the Shine-Dalgarno (SD) sequence of the mRNA (pink) with the anti-SD sequence of the 16S rRNA (tan). **f**, Superimposition of (**d**) and (**e**). **g**, Binding site of *E.coli* RMF (blue) on the hibernating 70S (E70S-RMF) compared with the (h) binding site *E. coli* RMF (cyan) on the *T. thermophilus* 70S (T70S-RMF), and (i) superimposition of (**g**) and (**h**). Ribosomal proteins S7 (salmon), S11 (green), S21 (orange) and the anti-SD sequence (tan) are shown for reference.

**Extended Data Figure 5.**
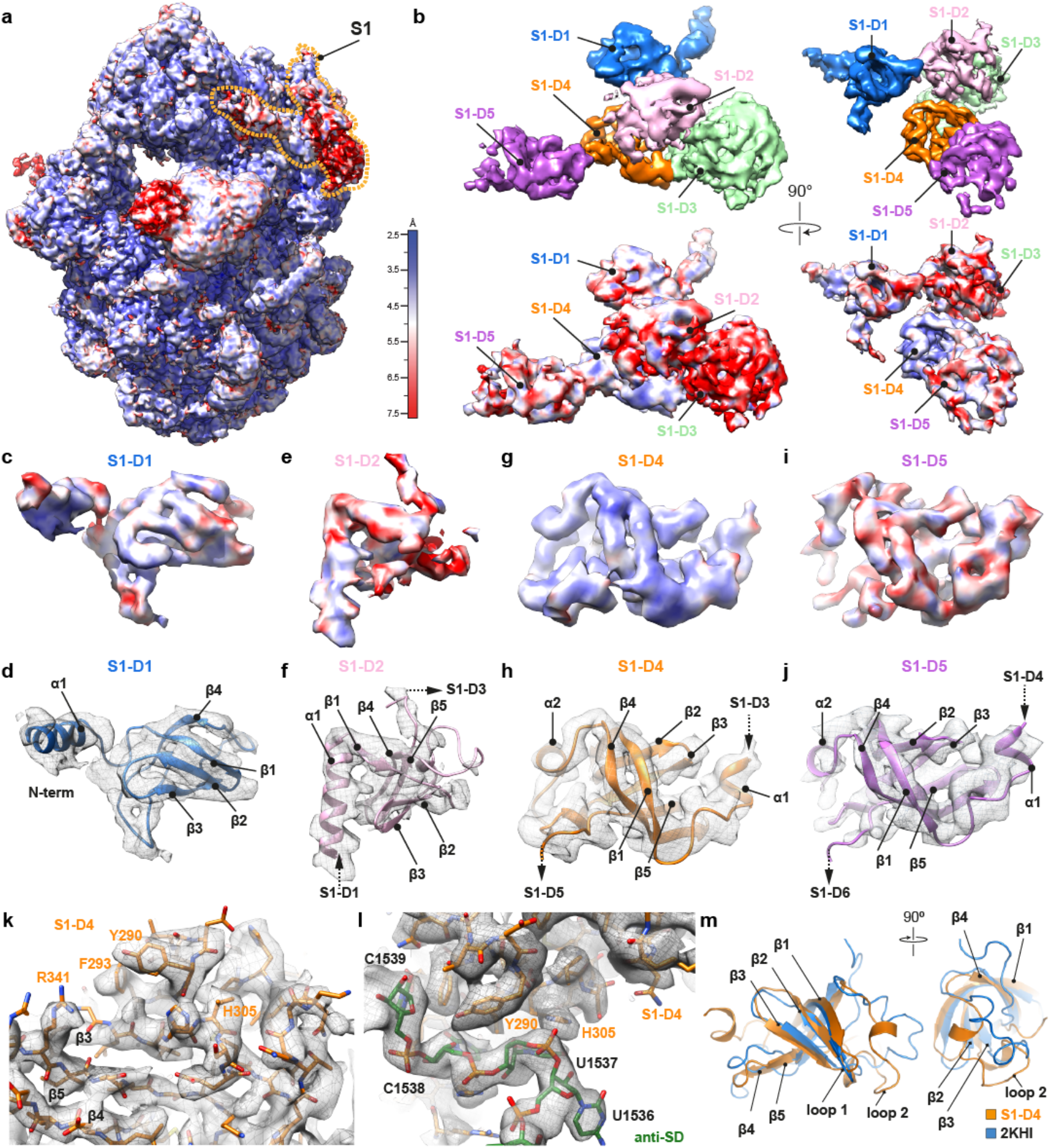
Molecular model for ribosomal protein S1. **a,** Overview of the hibernating 70S ribosome colored according to local resolution with binding position of S1 highlighted. **b,** Cryo-EM density colored by domain (upper panels) and by local resolution (lower panels). c-j, Isolated cryo-EM density for (**c-d**) S1-D1, (**e-f**) S1-D2, (**g-h**) S1-D4 and (**i-j**) D1-D5, (**c,e,g,i**) colored according to local resolution or (**d,f,h,j**) represented as mesh (grey) with fitted molecular models. The N- and C-termini that connect to neighboring domains are indicated with arrows. **k-l**, selected views showing cryo-EM density (grey mesh) with fitted models and sidechains for S1-D4 (orange) and nucleotides of anti-SD (green). **m**, The conformation of S1-D4 on the hibernating 70S ribosome (orange) compared with NMR structure of S1-D4 in solution (blue; PDB ID 2KHI)^23^.

**Extended Data Figure 6.**
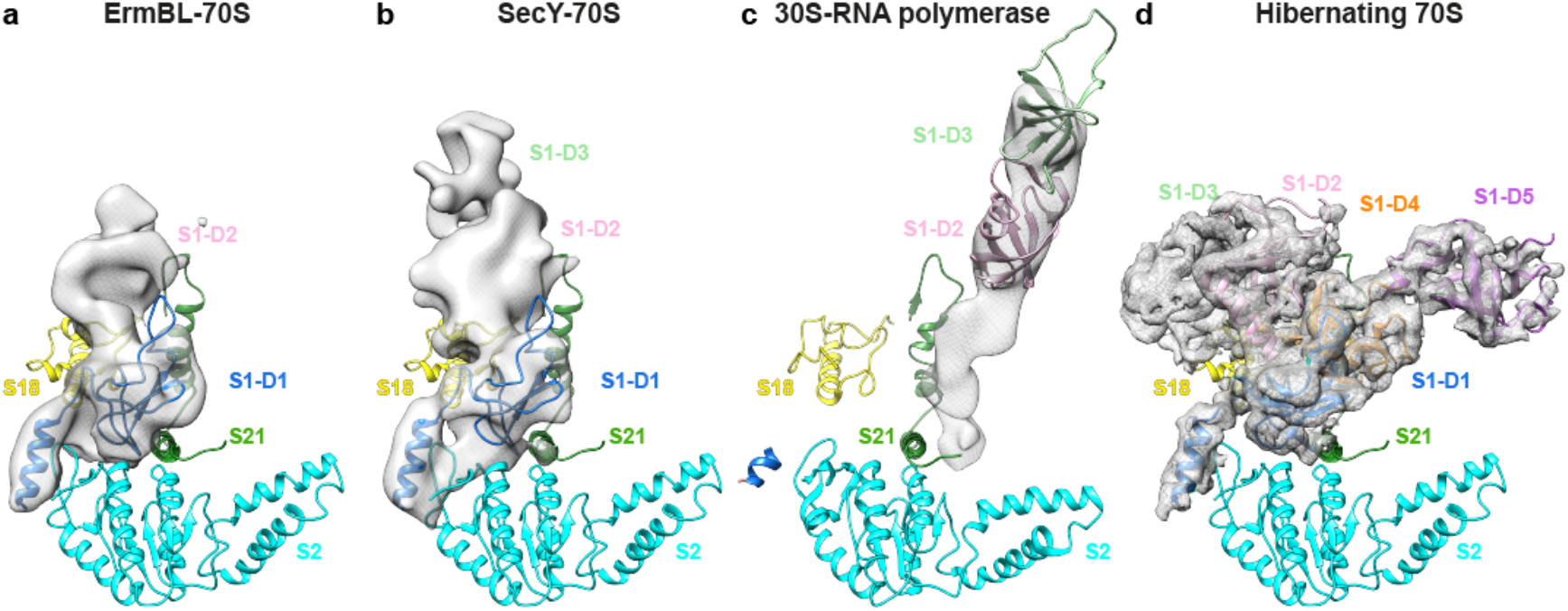
Conformation of S1 in various cryo-EM reconstructions. **a-d,** Cryo-EM density (grey) of S1 as observed in the (**a**) ErmBL-70S (EMD 6211)^21^, (**b**) SecY-70S (EMD 5693)^20^, (**c**) 30S-RNA polymerase complex (EMD 7014)^22^, (**d**) and on the hibernating 70S ribosome. Ribosomal proteins S2 (cyan), S18 (yellow) and S21 (green) are shown for reference.

**Extended Data Figure 7.**
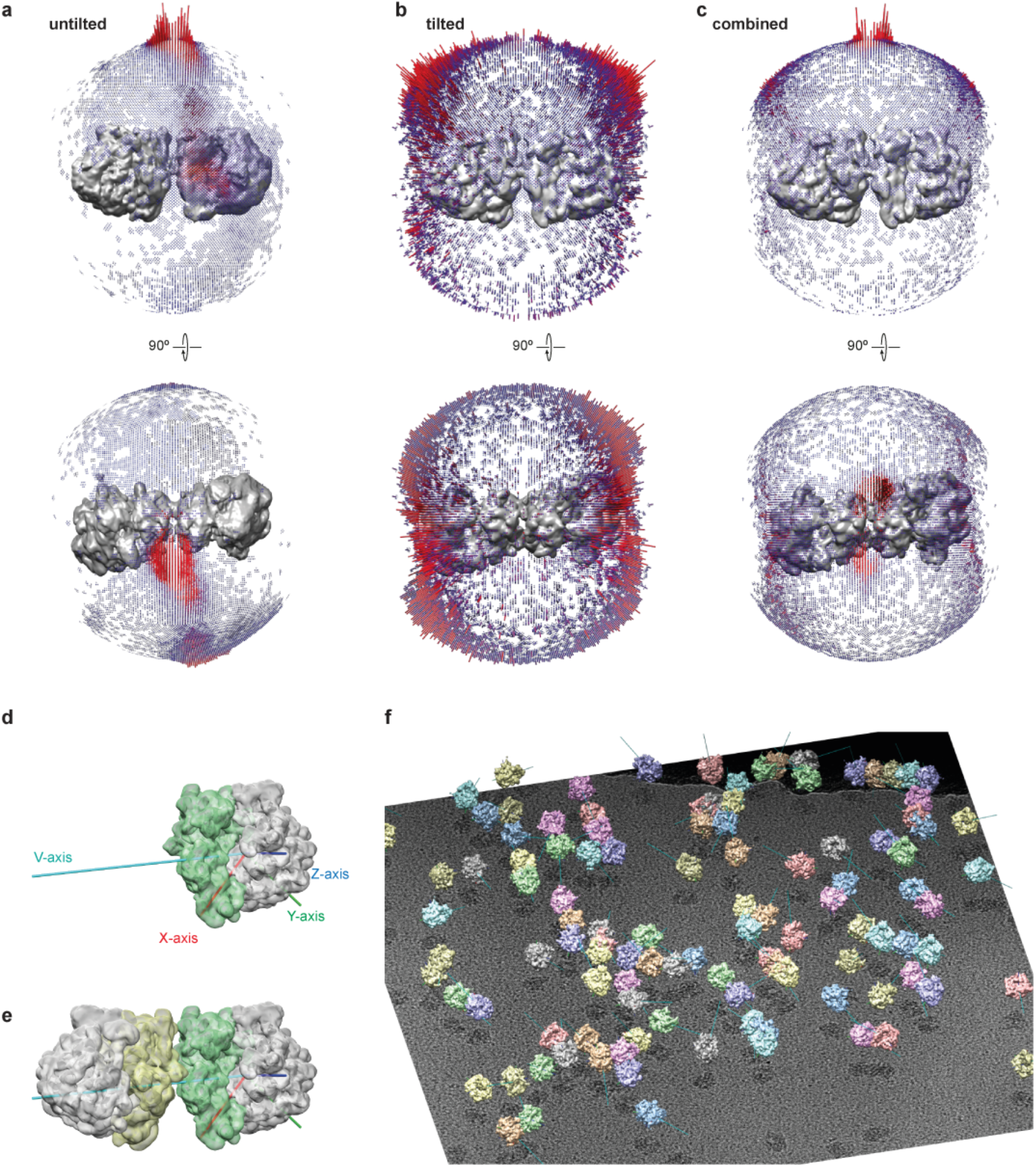
Data collection and 100S particle picking strategies. **a-c,** Euler angle distributions of the (**a**) untilted (0°), (**b**) 40° tilted and (**c**) combined (40° + 0°) data for the 100S particles. **d-e**, 70S and 100S images superimposed with the vectors used to pick 100S particles with the correct distance and relative orientation. f, 3D representation of the particles from one micrograph with the vectors used for particle picking displayed.

**Extended Data Figure 8.**
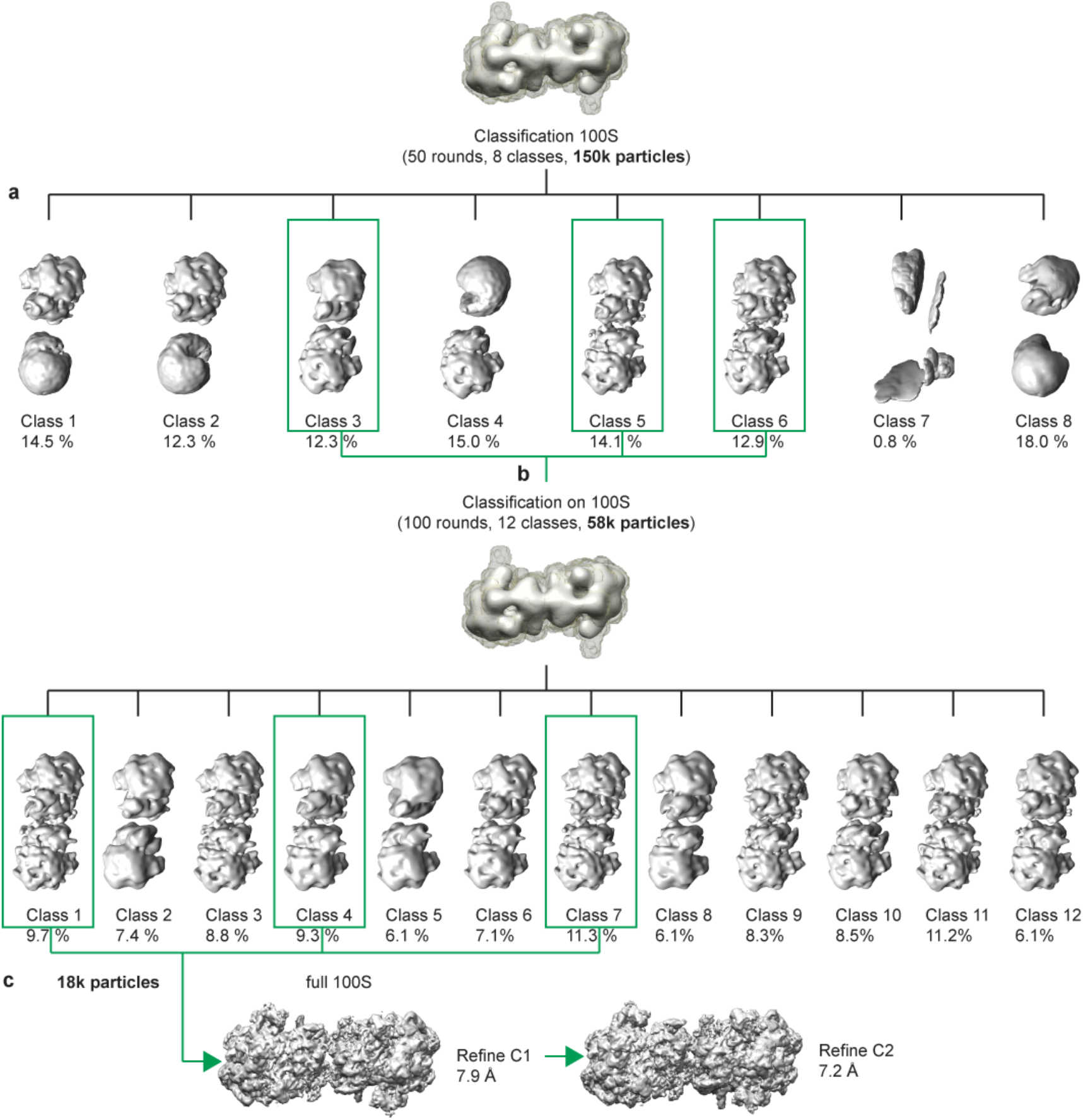
*In silico* sorting and processing of the 100S particle. **a,** Selected 100S (150,000 particles) were subjected to 50 rounds of 3D classification using a full 100S mask in relion, yielding 8 classes with varying degrees of stability. **b,** The three most stable classes (class 3, 5 and 6) were combined (58,000 particles total) and subjected to 50 further rounds of 3D classification. **c,** Again the three most stable classes (class 1, 4 and 7) were combined (18,000 particles total) and refined without symmetry yielding a cryo-EM structure of the 100S particle at 7.9 Å, or with C2 symmetry imposed yielding a cryo-EM structure of the 100S particle at 7.2 Å.

**Extended Data Figure 9.**
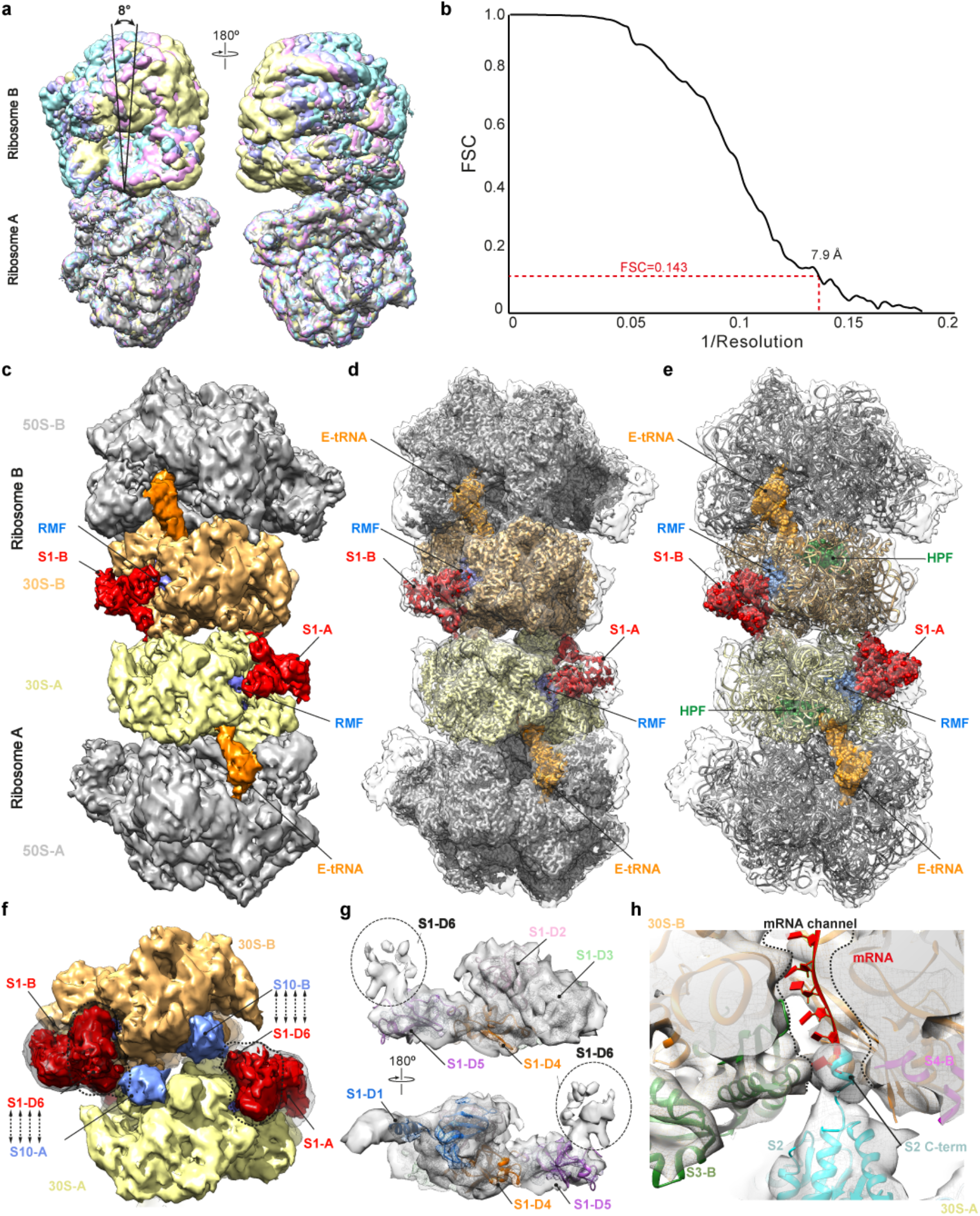
Cryo-EM structure of the hibernating 100S particle. **a,** Alignment of 100S particles from Class 1 (cyan), 2 (pink), 4 (purple) and 6 (yellow) based on ribosome A, revealing tilting of up to 8° of ribosome B with respect to Ribosome A. **b,** Fourier-shell correlation (FSC) curve of the final refined map, indicating that the average resolution of the hibernating 100S ribosome is 7.9 Å (at FSC 0.143). **c,** Cryo-EM map of the hibernating 100S particle, with isolated densities for 30S-A (yellow), 30S-B (tan), 50S-A/50S-B (grey), RMF (blue), HPF (green), E-tRNA (orange) and S1 (red). **d-e**, Rigid body fit of the cryo-EM map of the hibernating 100S particle (grey) with the (**d**) cryo-EM map of the hibernating 70S ribosome and (**e**) molecular model of the hibernating 70S ribosome. f, Cryo-EM map of the hibernating 100S particle with isolated densities for 30S-A (yellow) and S1-A/B (red) as well as 30S-B (tan) and S10-A/B (blue). **g,** Isolated density (grey mesh) for S1 protein from the hibernating 100S, fitted with molecular models for S1-D1 (blue), S1-D2 (salmon), S1-D3 (green), S1-D4 (orange) and S1-D5 (magenta). The region for S1-D6 is marked with an ellipse (black). **h,** Transverse section of the cryo-EM map (grey) of the hibernating 100S particle, revealing density for the C-terminal helix of S2-A (cyan) probing the mRNA entrance tunnel of 30S-B (16S rRNA in orange). Ribosomal proteins S3-B (green) and S4-B (magenta) that line the mRNA entrance tunnel are shown, as well as a superimposition of a mRNA (red).

**Extended Data Figure 10.**
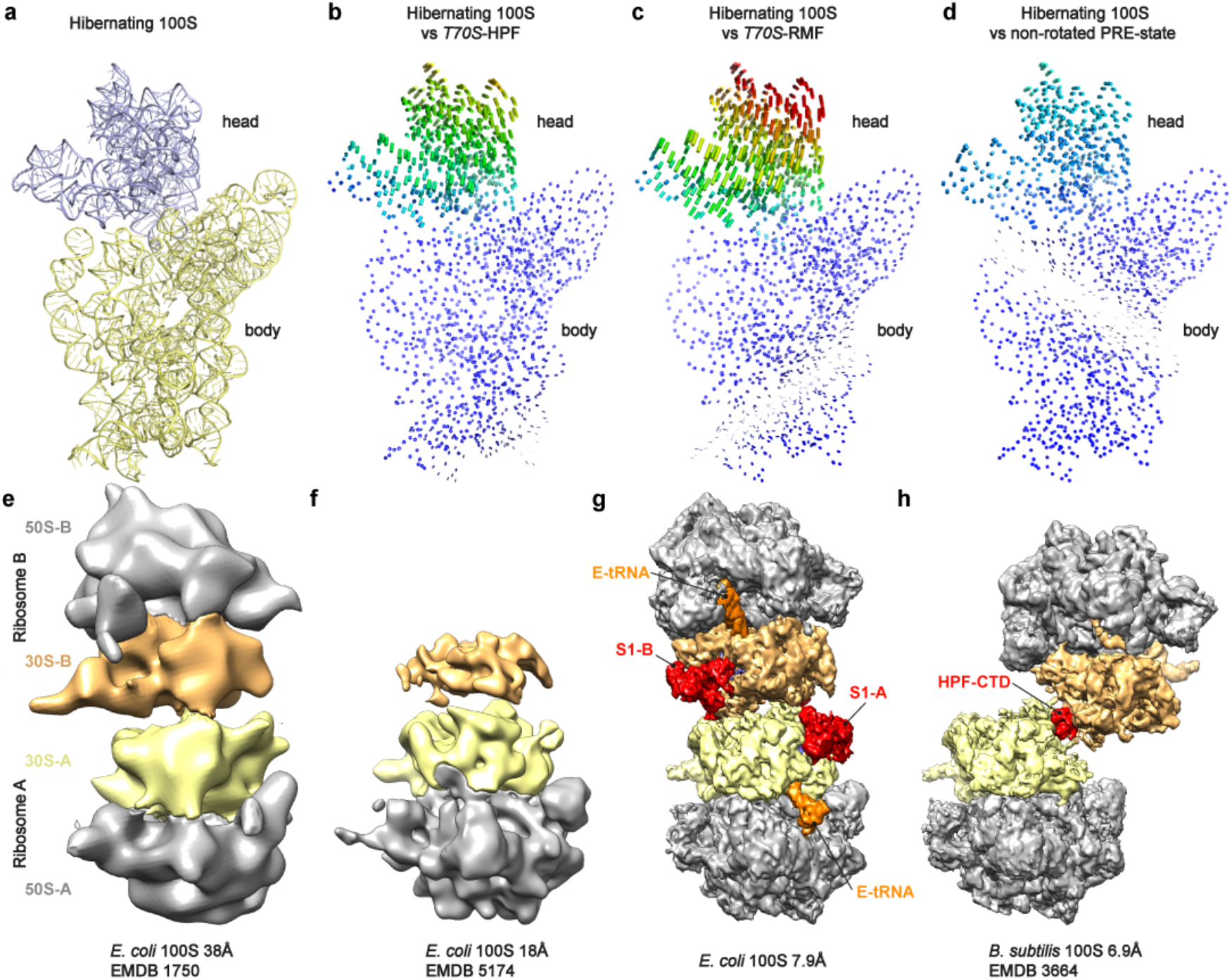
Conformation of the hibernating 100S particle. **a,** Model of the 16S rRNA of the 30S subunit of the hibernating 100S particle with head (slate) and body (yellow) highlighted. **b-c,** 30S subunit structures illustrating the degree of head swivel in the 30S subunit of the hibernating 100S particle relative to the structure of the (b) *E. coli* HPF in complex with the *T. thermophilus* 70S ribosome (T70S-HPF; PDB ID 4V8H)^13^, (c) *E. coli* RMF in complex with the *T. thermophilus* 70S ribosome (T70S-RMF; PDB ID 4V8G)^13^, and (**d**) non-rotated pre-translocation (PRE) state ribosome (PDB ID 1VY4)^48^. The distance each atom shifts relative to the reference structure is directly colored on the small subunit, as shown by colored lines connecting the same atoms between the reference and the shifted structure. **e-h,** comparison of the cryo-EM maps of the *E. coli* 100S particle at (**e**) 38 Å (EMD 1750)^6^, (**f**) 18 Å (EMD 5174)^5^ and (g) determined in this study as 7.9 Å, relative to (**h**) the cryo-EM of a Gram-positive 100S particle (EMD 3664)^7^.

**Extended Data Figure 11.**
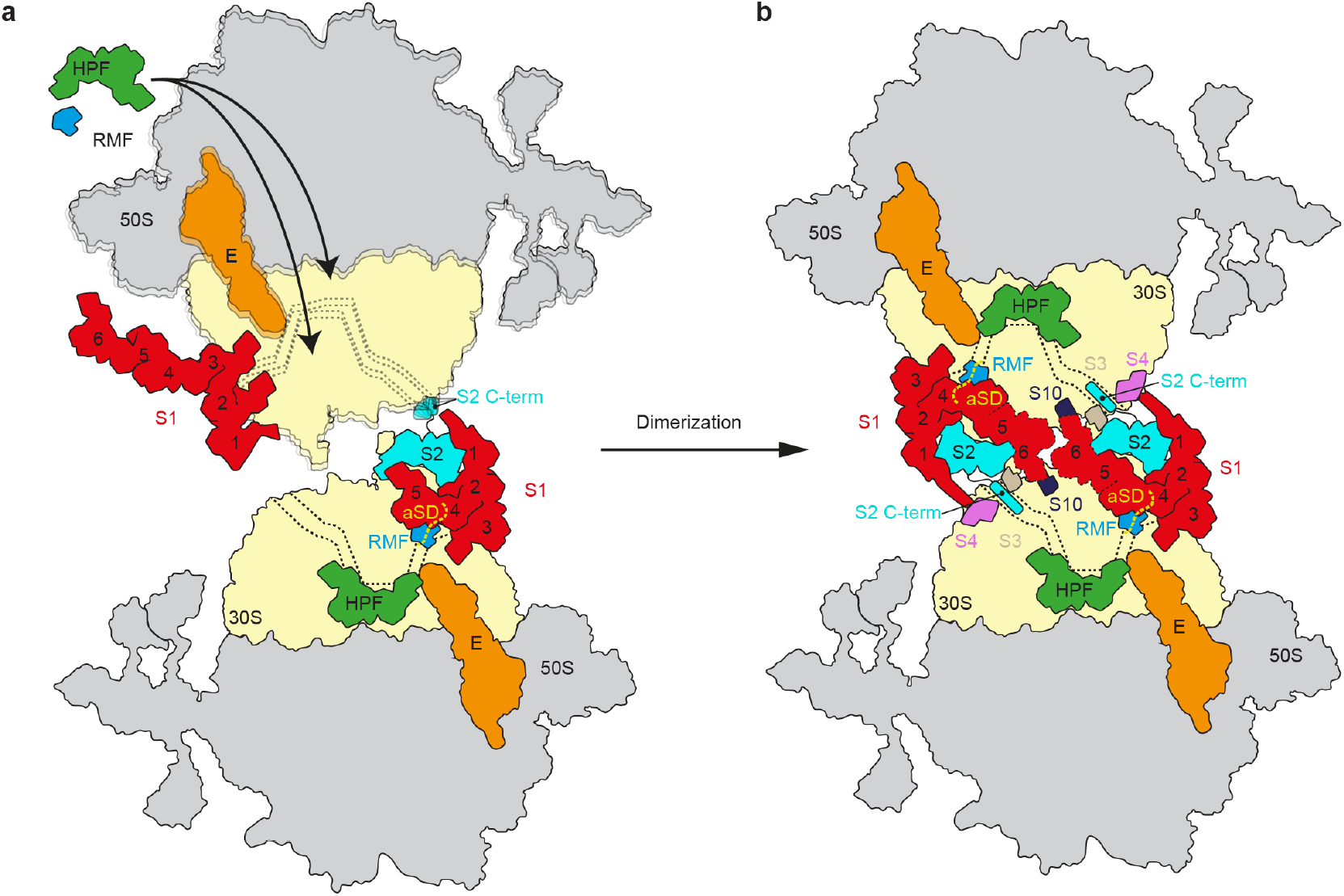
Model for RMF and HPF mediated 100S formation. **a,** Binding of HPF, RMF and E-tRNA to vacant 70S ribosomes leads to stabilization of a non-rotated ribosome with a defined non-swiveled head conformation. The presence of HPF blocks the binding of mRNA and tRNAs to the A- and P-sites and stabilizes the E-tRNA. RMF binding stabilizes a compacted conformation of S1 where domains 4-6 fold back into the 30S subunit rather than extending into the solvent. **b,** Dimerization is facilitated by the compacted conformation of S1, promoting interactions between the N-terminus of S1-D1 with S4 and S1-D6 with S10. In addition, the C-terminus of S2 probes the mRNA entry channel of the neighboring ribosome. Inactivation of the 100S particle is also facilitated by RMF and S1 that directly interact with the anti-Shine-Dalgarno sequence of the 16S rRNA, preventing recruitment of mRNA to the ribosome during translation initiation.

Cryo-EM data collection, refinement and validation statistics.

**Cryo-EM data collection, refinement and validation statistics**
|  | Hibernating 70S<br>(EMDB-2XYZ)<br>(PDB 4XYZ) | 100S particle<br>(EMDB-3XYZ)<br>(PDB 5XYZ) |
| --- | --- | --- |
| **Data collection and processing** |  |  |
| Magnification | 129,151 | 105,000 |
| Voltage (kV) | 300 | 300 |
| Electron exposure (e<sup>-</sup>/Å<sup>2</sup>) | 28 | 51 |
| Defocus range (μm) | -0.8 to -2.5 | -1.2 to -3.5 |
| Pixel size (Å) | 1.084 | 1.35 |
| Symmetry imposed | C1 | C1,C2 |
| Initial particle images (no.) | 625,839 | 349,000 |
| Final particle images (no.) | 188,304 | 18,000 |
| Map resolution (Å) | 3.0 | 7.9 |
| FSC threshold | 0.143 | 0.143 |
| Map resolution range (Å) | 2.5-7.5 |  |
| **Refinement** |  |  |
| Initial model used (PDB code) | 5O2R |  |
| Model resolution range (Å) | 3.2 |  |
| Map sharpening *B* factor (Å<sup>2</sup>) | -72.5 |  |
| Correlation coef (%; Phenix) | 0.7665 |  |
| Model composition |  |  |
| Non-hydrogen atoms | 145,806 |  |
| Protein residues | 5,926 |  |
| RNA bases | 4,636 |  |
| *B* factors (Å<sup>2</sup>) |  |  |
| Full Complex | 44.2 |  |
| R.m.s. deviations |  |  |
| Bond lengths (Å) | 0.007 |  |
| Bond angles (°) | 1.02 |  |
| Validation |  |  |
| MolProbity score | 1.84 |  |
| Clashscore | 5.41 |  |
| Poor rotamers (%) | 0.62 |  |
| Ramachandran plot |  |  |
| Favored (%) | 90.14 |  |
| Allowed (%) | 9.57 |  |
| Disallowed (%) | 0.29 |  |
| Validation RNA |  |  |
| Correct sugar pucker (%) | 98.5 |  |
| Good backbone conf. (%) | 97.9 |  |

